# Laser microdissection, proteomics and multiplex immunohistochemistry: a bumpy ride into the study of paraffin-embedded fetal and pediatric lung tissues

**DOI:** 10.1101/2023.07.25.550475

**Authors:** L.M. Cardoso dos Santos, Y. Avila, D. Schvartz, A-L Rougemont, M-L Bochaton-Piallat, I. Ruchonnet-Metrailler

**Author notes:** **Correspondence:** I. Ruchonnet-Metrailler.

## Abstract

**Background:** Knowledge about lung development or lung disease is mainly derived from data extrapolated from mouse models. This comes with obvious drawbacks in developmental diseases, particularly due to species differences. Our objective is to describe the development of complementary analysis methods that will allow a better understanding of the molecular mechanisms involved in the pathogenesis of rare congenital diseases.

**Methods:** Paraffin-embedded human pediatric and fetal lung samples were laser microdissected to enrich different lung regions, namely bronchioli or alveoli. These samples were analyzed by data independent acquisition-based quantitative proteomics and lung structures were subsequently compared. To confirm the proteomic data, we employed and optimized <u>S</u>equential <u>IM</u>muno<u>P</u>eroxidase <u>L</u>abeling and <u>E</u>rasing (SIMPLE) staining for specific proteins of interest.

**Results:** By quantitative proteomics, we identified typical pulmonary proteins from being differentially expressed in the different regions. While the receptor for advanced glycation end products (RAGE), surfactant protein C (SFTPC) were downregulated, tubulin beta 4B (TUBB4B) was upregulated in bronchioli, compared to alveoli. In fetal tissue, CD31 was downregulated in fetal bronchioli, compared to canaliculi. Moreover, we confirmed their presence using SIMPLE staining. Some expected proteins did not show up in the proteomic data, like SOX-9 that was only detected by means of immunohistochemistry in the SIMPLE analysis.

**Conclusion:** Our data underlines the robustness and applicability of this type of experimental approach, especially for rare paraffin-embedded tissue samples. It also strengthens the importance of these methods for future studies, in particular, when considering developmental lung diseases, such as congenital lung anomalies.

## 1 Introduction

The lung is an essential organ to the life of all mammals. Nonetheless, there is still a big gap in the knowledge about its development due to the complexity of the physiology and biological mechanisms occurring in this organ. Consequently, the lack of *in vivo* and *in vitro* models, hinders the proper understanding of various pathologies, and in particular of rare developmental congenital anomalies. To overcome this issue, researchers have been developing new techniques that are able to provide more detailed information using the scarce material that is available from patients, typically formalin-fixed paraffin-embedded (FFPE) tissues. Many biobanks contain a multiplicity of these valuable samples that can be used to further study these pathologies in a retrospective manner. This type of preservation is particularly attractive due to the convenience in handling the material and its excellent tissue morphology preservation. For protein detection in tissue samples, both in a clinical diagnostic setting and for research issues, immunohistochemistry (IHC) is the gold standard; however, unless multiplexing is used, it does not allow the co-visualization of multiple targets in one section, while tissue availability may become an issue in small biopsy specimens. Immunofluorescence staining came as an alternative, but generally at the cost of diminished data quality, e.g. autofluorescence (particularly present in lung tissue), photobleaching when acquisitioning big areas of tissue, limited number of fluorophore combinations, and often the need of different antibodies than those already used in histopathology laboratories.

Herein, we propose a coupled approach to study in-depth the proteome of human lung, using laser microdissected (LMD) FFPE tissue samples, associated with a multiplexed IHC technique, coined <u>S</u>equential <u>IM</u>muno<u>P</u>eroxidase <u>L</u>abeling and <u>E</u>rasing (SIMPLE). The latter will confirm the proteomic data, while providing the localization of several target proteins in one tissue section. The LMD technique allows to define and isolate specific histological structures in FFPE sections with enrichment of a defined cellular or tissular compartment, overcoming tissue heterogeneity (1-3). Proteomic studies, particularly LMD-associated proteomics, have been used in the characterization of the human lung. It has been employed to characterize normal lung tissue in FFPE samples, focusing on the extracellular matrix composition (4), or define the lung postnatal developmental protein signature (5). On a pathological context, we previously performed LMD proteomics on congenital pulmonary airway malformation human samples (6). This previous work provided only qualitative data, due to lack of material, a situation that is currently being improved in a follow-up study. The confirmation of proteomic data using the SIMPLE technique has already been described for different organs (7-9) but, to the best of our knowledge, never in pediatric and fetal lung tissue.

## 2 Materials and Methods

### 2.1 Tissue sample preparation and laser microdissection (LMD)

FFPE blocks of human pulmonary tissue specimens from 5 fetuses (Ethical committee 2016-00175) and 5 healthy lung tissue of postnatal pulmonary lobectomies (Ethical committee 12-110) (**Supplementary table 1**) were cut into 10 μm-thick sections and mounted on Leica PET-membrane slides (76463-320, Leica Microsystems). Postnatal lung tissues were considered as “healthy” after microscopic/histological analysis by a pulmonary pathologist. Samples were deparaffinized and hydrated according to standard histological procedures, stained with hematoxylin (S330130-2, Agilent) and stored at 4 °C. LMD was performed in all 10 hematoxylin-stained samples (**Figure 1A**), according to manufacturer user guidelines. Briefly, an upright Leica DM6500 microscope, equipped with a Cryslas laser and a Leica CC7000 camera, was used to perform the tissue LMD, where specific tissue fragments were isolated with a UVI 5L×/0.12 microdissection objective and regions of interest were defined manually with the LAS-AF software (Leica). The isolated fragments were collected in 0.5 ml PCR tubes. Briefly, we proceeded into an enrichment in the epithelial portions of postnatal and fetal lung tissues from different lung structures: postnatal bronchioli and alveoli, and fetal bronchioli and canaliculi, respectively. Noteworthy, due to the nature of the technique, the enrichment in the epithelial component was not complete, including as well the immediate surrounding mesenchyme, namely the bronchiolar muscle layer and capillaries. Structures were selected based on their histological features. Bronchioli, both fetal and postnatal, were defined as well-differentiated airways lined by a pseudo-stratified columnar epithelium or a simple cuboidal epithelium supported by smooth muscle and connective tissue. In fetal lung at the canalicular stage of development, canaliculi were defined as tubular structures varying from simple to branched, lined by columnar vacuolated epithelium to cuboidal non-vacuolated epithelium. Alveoli were easily recognized, and care was taken not to include medium-sized blood vessels or lymphoid aggregates.

**Figure 1.**
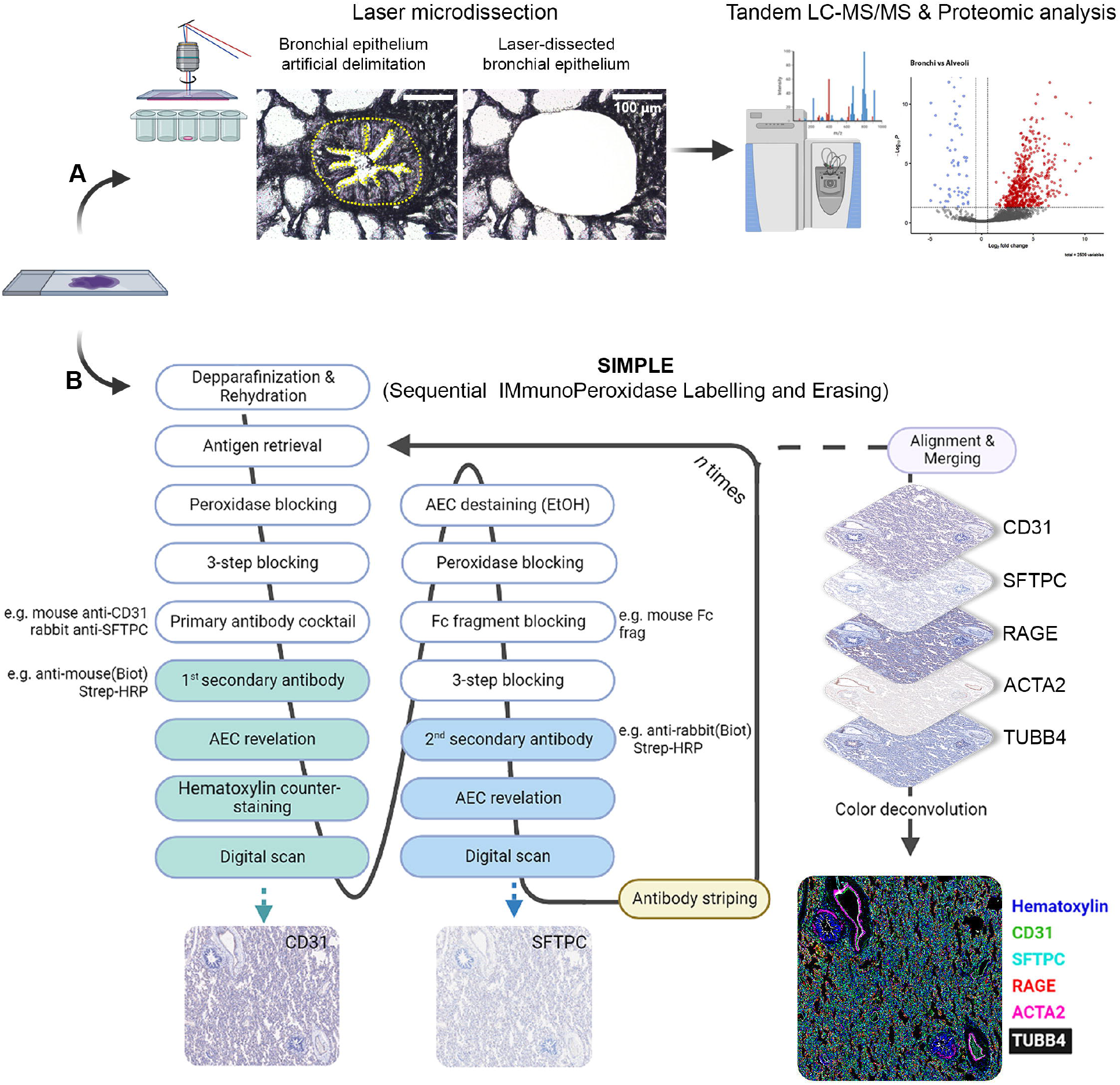
Scheme depicting LMD proteomic approach and SIMPLE stammg. **(A)** Formalin-fixed paraffin-embedded (FFPE) samples of fetal and postnatal bronchioli (represented in the figure), canaliculi and alveoli were laser microdissected to enrich their epithelial portions, and subsequently subjected to liquid chromatography coupled with tadem mass spectrometry (LC-MS/MS) and proteomic analysis. **(B)** FFPE samples of fetal and postnatal lung tissue were stained following the SIMPLE protocol. Briefly, samples are depparafinized and rehydrated, entering a cycle of subsequent stainings. At each cycle, samples go through a step of antigen retrieval and 4 steps of blocking, followed by an incubation with a primary antibody cocktail (note that the antibodies should be from different species). Each of the secondary antibodies is revealed sequentially, digital images are acquired and AEC staining removed by ethanol washing. When the first cycle is finished, the staining process re-starts by the antigen retrieval step to remove the primary antibodies. After the entire staining process is concluded, brightfield images are aligned, merged and artificially colored.

Samples were collected and stored at -20 °C. To normalize tissue quantity for subsequent mass spectrometry (MS)-based proteomics, the equivalent of 1.2 × 10^6^ μm^2^ for the bronchioli, fetal bronchioli and canaliculi area, and of 3 × 10^6^ μm^2^ for the alveoli area were collected. The procedure was performed as described by Drummond and colleagues (3), who obtained a good compromise between proteomic data retrieval and hands-on microdissection time. The difference between the LMD alveoli surface and other tissues was calculated beforehand in order to compensate its lower cellularity per μm^2^, typical of the honeycomb-like structure of pulmonary parenchyma.

### 2.2 Proteomic sample preparation

When required, LMD samples (n=5 for each sample structure) were pooled by adding 60 μl of acetonitrile (10055454, Thermo Fisher Scientific) to each tube and spun down, then pooled in a new 1.5 mL Eppendorf tube and dried under speed-vacuum. Samples were resuspended in 0.1% RapiGest Surfactant (186001860, Waters) in 50 mM ammonium bicarbonate as to obtain a ratio of 1.2 × 106 μm^2^ per 100 μl. They were sonicated (6 × 30 s) at 70% amplitude and 0.5 pulse and kept on ice 30 s between each cycle of sonication. To remove residual paraffin and sufficiently reverse formaldehyde crosslinks samples were heated for 1 h at 95 °C, then 1 h at 65 °C, and followed by a second round of sonication. For 100 μl of sample, 2 μl of 50 mM dithioerythritol were then added and the reduction was carried out at 37 °C for 1 h. Alkylation was performed by adding 2 μl of 400 mM iodoacetamide (per 100 μl of sample) during 1 h at room temperature in the dark. Overnight digestion was performed at 37 °C with 2 μL of freshly prepared trypsin at 0.1 μg/μl (V5111, Promega) in 50 mM ammonium bicarbonate (per 100 μl of sample). To remove RapiGest, samples were acidified with 5 μl of 10% trifluoroacetic acid (91719, Sigma-Aldrich), heated at 37 °C for 45 min and centrifuged 10 min at 17’000 x g. Each sample was desalted with a C18 microspin column (Harvard Apparatus) according to manufacturer’s instructions, completely dried under speed-vacuum and stored at -20°C.

### 2.3 Liquid chromatography electrospray ionization tandem mass spectrometry (LC/ESI-MS/MS)

Samples were resuspended in 12 μl of loading buffer (5% CH_3_CN, 0.1% CH_2_O_2_). iRT peptides (Biognosys) were added to each sample and 4 μl of peptides were injected on column. LC/ESI-MS/MS was performed on an Orbitrap Fusion Lumos Tribrid mass spectrometer (Thermo Fisher Scientific) equipped with an Easy nLC1200 liquid chromatography system (Thermo Fisher Scientific). Peptides were trapped on an Acclaim pepmap100, C18, 3 μm, 75 μm x 20 mm nano trap-column (Thermo Fisher Scientific) and separated on a 75 μm x 500 mm, C18 ReproSil-Pur (Dr. Maisch GmBH), 1.9 μm, 100 Å, home-made column. The analytical separation was run for 135 min using a gradient of H_2_O/CH_2_O_2_ (99.9%/0.1% - solvent A) and CH_3_CN/H_2_O/CH_2_O_2_ (80.0%/19.9%/0.1% - solvent B). The gradient was run from 8% to 28 % of solvent B in 110 min, to 42% of solvent B in 25 min, then to 95% solvent B in 5 min, with a final stabilization step of 20 min in 95% solvent B. Flow rate was of 250 nL/min, with a total run time of 160 min. Data-Independant Acquisition (DIA) was performed with a precursor full scan (MS1) at a resolution of 60’000 full width at half maximum (FWHM), followed by 30 product ion scans (MS2) with fix wide precursor isolation windows. MS1 was performed in the Orbitrap with an AGC target of 1 × 10^6^, a maximum injection time of 50 ms and a scan range from 400 to 1240 m/z. DIA MS2 was performed in the Orbitrap using higher-energy collisional dissociation (HCD) at 30%. Isolation windows was set to 28 m/z with an AGC target of 1 × 10^6^ and a maximum injection time of 54 ms. Due to the typically low yield of protein, we performed a single run of LC-MS/MS in 5 different biological samples for each structure type.

### 2.4 Proteomic data analysis

DIA raw files were loaded into Spectronaut v.15 (Biognosys) and analyzed by directDIA using default settings. Briefly, data were searched against human reference proteome database (Uniprot; 20660 entries). Trypsin was selected as the enzyme, with one potential missed cleavage. Variable amino acid modifications were oxidized methionine and deaminated (NQ). Fixed amino acid modification was carbamidomethyl cysteine. Both peptide precursor and protein false discovery rate (FDR) were controlled at 1% (*Q value* < 0.01). Single hit proteins were excluded. For quantitation, Top 3 precursor area per peptides were used, “only protein group specific” was selected as proteotypicity filter and normalization was set to “global normalization”. The quantitative analysis was performed with MapDIA tool, using the precursor quantities extracted from Spectronaut output. No further normalization was applied. The following parameters were used: min peptides = 2, max peptides = 10, min correl = -1, Min_DE = 0.01, max_DE = 0.99, and experimental_design = Independant design. For subsequent analysis, each group (postnatal and fetal bronchioli, alveoli and canaliculi) were composed by the 5 different biological samples. Proteins were considered to have significantly changed in abundance with an FDR ≤ 0.05 and an absolute fold change FC≥ |1.5| (log2FC ≥ |0.58|). Gene Ontologies term enrichment analysis, as well as associated plots, were performed with ClusterProfiler (10, 11) R package, using the EnrichGO function. Enrichment tests were calculated for GOterms, based on hypergeometric distribution. Enrichment p-value cutoff was set to 0.05, and Q-value cutoff to 0.01. The gofilter function was use prior to cnetplot drawing, to filter results at specific levels. To determine protein cellular localization, we used the NeXtProt knowledge database (reviewed and annotated for the human proteome) and extracted the gold and silver annotated proteins for four Gene Ontologies and Uniprot Subcellular Location (SL) terms: plasma membrane (GO:0005886 and SL-0036), extracellular matrix (GO:0031012), cytosol (GO:0005829 and SL-0091) and nuclear (GO:0005634 and SL-0191) (**Supplementary table 2**). The mass spectrometry proteomics data (DIA raw files) have been deposited to the ProteomeXchange Consortium via the PRIDE (12) partner repository with the dataset identifier PXD040654.

### 2.5 Sequential IMmunoPeroxidase Labeling and Erasing (SIMPLE)

A schematic representation of the SIMPLE staining is depicted in **Figure 1B**. Three μm FFPE tissue sections were mounted on TruBond™ 380 slides (50-364-505, Electron Microscopy Sciences), placed at 54°C overnight before going through deparaffination, 4 × 10 min in Ultraclear (3905.9010PE, Avantor), and hydrated in a series of graded alcohols (95 - 50%) to distilled water. Specimen went through heat-induced epitope retrieval (HIER) in either citrate pH 6.0 or tris-EDTA pH 9.0 buffers, according to antibody specification, using a Bio SB TintoRetriever Pressure Cooker (BSB-7087, Bio SB), set at low pressure for 10 min. After HIER, slides were cooled down in an ice bath for 15 min, endogenous peroxidases were quenched with DAKO REAL Peroxidase-Blocking Solution (S202386-2, Agilent) for 5 min. In addition to blocking unspecific antibody binding using Serum-Free Protein Block (X0909, Agilent) for 10 minutes, an Avidin/Biotin Blocking Kit (ab64212, Abcam) was used for 10 min to block endogenous avidin and biotin binding sites, respectively. Primary antibodies (**Supplementary table 3**) were then incubated for 1 h at room temperature.

The following antibodies were used: anti-CD31 (66065-2-lg, Proteintech), anti-surfactant protein C (SFTPC, AB3786, Merck), homemade anti-α-smooth muscle actin (clone 1A4, α-SMA or ACTA2 (13)) anti-receptor for advanced glycation end products (RAGE, ab3611, Abcam); anti-tubulin beta 4B (TUBB4B, sc-134230, Santa Cruz), and anti-SOX-9 (AB5535, Millipore). As negative controls, mouse IgG1 (BE0083, Bioxcell) or IgG2a (MAB0031, R&D Systems) isotype control, and rabbit IgG isotype control (10500C, Invitrogen) were used. Goat anti-mouse IgG (1:250, 115-066-146, Jackson ImmunoResearch) and goat anti-rabbit IgG (1:250, 111-066-144, Jackson ImmunoResearch) secondary biotinylated antibodies were incubated 30 min at room temperature, followed by another 30 min room temperature incubation with streptavidin/horseradish peroxidase (1:300, P0397, Agilent). Staining was obtained by using an alcohol-soluble peroxidase substrate, 3-amino-9-ethylcarbazole (AEC, AC1331003, DCS Innovative Diagnostik-Systeme). Slides were counterstained with hematoxylin and finally cover slipped with Aquatex mounting solution (1.08562.0050, Merck) or PBS before whole-slide image scanning at 20x magnification using a Zeiss AxioScan.Z1.

After image acquisition, slides were de-coverslipped by immersion in dH_2_O. AEC staining was then removed by sequential washing in 50% (1 min)/75% (3 min)/95% (8 min, shaking every 2 min)/75% (2 min)/50% (1 min) ethanol solutions, followed by rehydration in dH_2_O. Antigen/antibody complex stripping was achieved via HIER (10 min at high pressure). Mouse (1:25, 115-007-003; Jackson ImmunoResearch) and rabbit (1:25, 111-007-003, Jackson ImmunoResearch) fragment antigen-binding solutions targeted against primary antibodies host species were used to prevent undesired binding between not fully stripped antigen-antibody complex from the previous stain and the secondary biotinylated antibodies of the subsequent staining. At this point, slides re-enter the same process of staining as described above, starting with the Serum-Free Protein Block. All antibodies and Streptavidin/HRP were diluted in DAKO Antibody Diluent (S0809, Agilent).

### 2.6 SIMPLE data processing

Whole-slide multiplexed images originated from the same tissue section and consisting of 5 antigen revelation were processed with a custom-made framework developed in Matlab R2022b (The MathWorks). Briefly, these large data (tens of gigabyte) were aligned, at a cellular level, according to the same reference image using manual control points pairing to infer affine transformations. Afterward, hematoxylin and AEC chromogenic stainings of each image were unmixed using an adaptation of the work of Landini and colleagues (14) optimized for computation speed and processing of large dataset. Finally, a single multiplexed image, with one channel per antigen and the related metadata, was exported in the OME-TIFF file format. This newly generated images could then be opened and processed with QuPath0.3.2 (15).

## 3 Results

### 3.1 Laser microdissection and proteomic analysis

For proteomic purposes, we collected 5 samples from either postnatal or fetal lung tissue (**Supplementary table 1**) that were laser-microdissected, enriching the epithelial portions of bronchioli, fetal bronchioli, alveoli and canaliculi structures (**Figure 2A and 2B**). Noteworthy, due to the structure of alveoli and canaliculi, these compartments correspond not only to the alveoli and canaliculi but also to the immediate surrounding mesenchyme. For each individual, several tissue fragments were microdissected, as shown in (**Figure 2A and 2B**) where the minimum amount of selected area (SA) tissue was calculated as to include the laser-destroyed area (LA) and provide enough material to allow downstream proteomic analysis. Principal component analysis (PCA) confirmed that LMD is technically reproducible and, most importantly, that samples from the same experimental group are in general close in their protein profile, thus strengthening the proteomics analysis (**Supplementary figure 1**). The main differences in protein pattern were found in PC1 between alveoli and fetal bronchioli (43% of the model variability). Moreover, we observed that the alveolar samples were the only group that presented considerable variability. As well, one of the samples from the postnatal bronchioli was a clearly apart from the rest of the samples in the group (**Supplementary figure 1**, sample E5), which we decided to maintain for discussion purposes. Following, we focused on the proteomic differences between bronchioli and alveoli (**Figure 2C**), or their fetal counterparts, fetal bronchioli and canaliculi (**Figure 2D**). For the two comparisons we detected in total 2509 and 2548 proteins, respectively, which were then assigned a cellular localization. The proportion of the assigned localizations were similar in the two sets of data, roughly corresponding to 60% nuclear, 47% cytoplasmic, 30% plasma membrane or 6% extracellular matrix (**Supplementary figure 2, Supplementary table 2**). Of note, in both comparisons, 44% of all proteins share at least 2 cellular localizations. For the purpose of this study, among the 592 and 484 proteins that were found differentially expressed (FDR < 0.05; Fold change [Log2] ≥ 0.58) in the comparison of bronchioli vs alveoli and fetal bronchioli vs canaliculi, respectively, we focused on proteins that are characteristic of bronchial and alveolar structures. For the bronchial compartment we selected TUBB4B and ACTA2, markers of ciliary epithelium and airway smooth muscle cells, respectively. As for the alveolar structures, we selected the RAGE, SFTPC and CD31, as markers of alveolar type I cells, alveolar type II cells, and endothelial cells, respectively. In postnatal tissue, TUBB4B and ACTA2 were upregulated in bronchioli, compared to alveoli. As expected, we found RAGE, SFTPC and CD31 significantly downregulated in bronchioli, compared to alveoli (**Figure 2Ca**). Regarding the fetal tissues (canalicular developmental stage), RAGE and SFTPC were considered not present or below the threshold of detection, not allowing the subsequent processing and quantification. Moreover, while TUBB4B expression was increased in fetal bronchioli, CD31 expression was increased in canaliculi (**Figure 2Da**). Following the differential protein expression analysis, we performed gene ontology enrichment of biological processes (BP) for postnatal and fetal samples (**Supplementary table 4**). For visualization purposes, the graphic representations of the enriched BP of the comparison “bronchioli vs alveoli” (**Figure 2Cb)** and “fetal bronchioli vs canaliculi” (**Figure 2Db**) did not include all branches of the GO tree. Five selected GO terms enriched in bronchiole compared to alveoli were grouped in three main classes related to muscle structure development, intracellular transport and mRNA maturation (**Figure 2Cb**). Similarly, in the fetal tissue, BP enriched in fetal bronchioli compared to canaliculi were grouped in three main classes related to tissue development, intracellular movement and cell movement (**Figure 2Db**).

**Figure 2.**
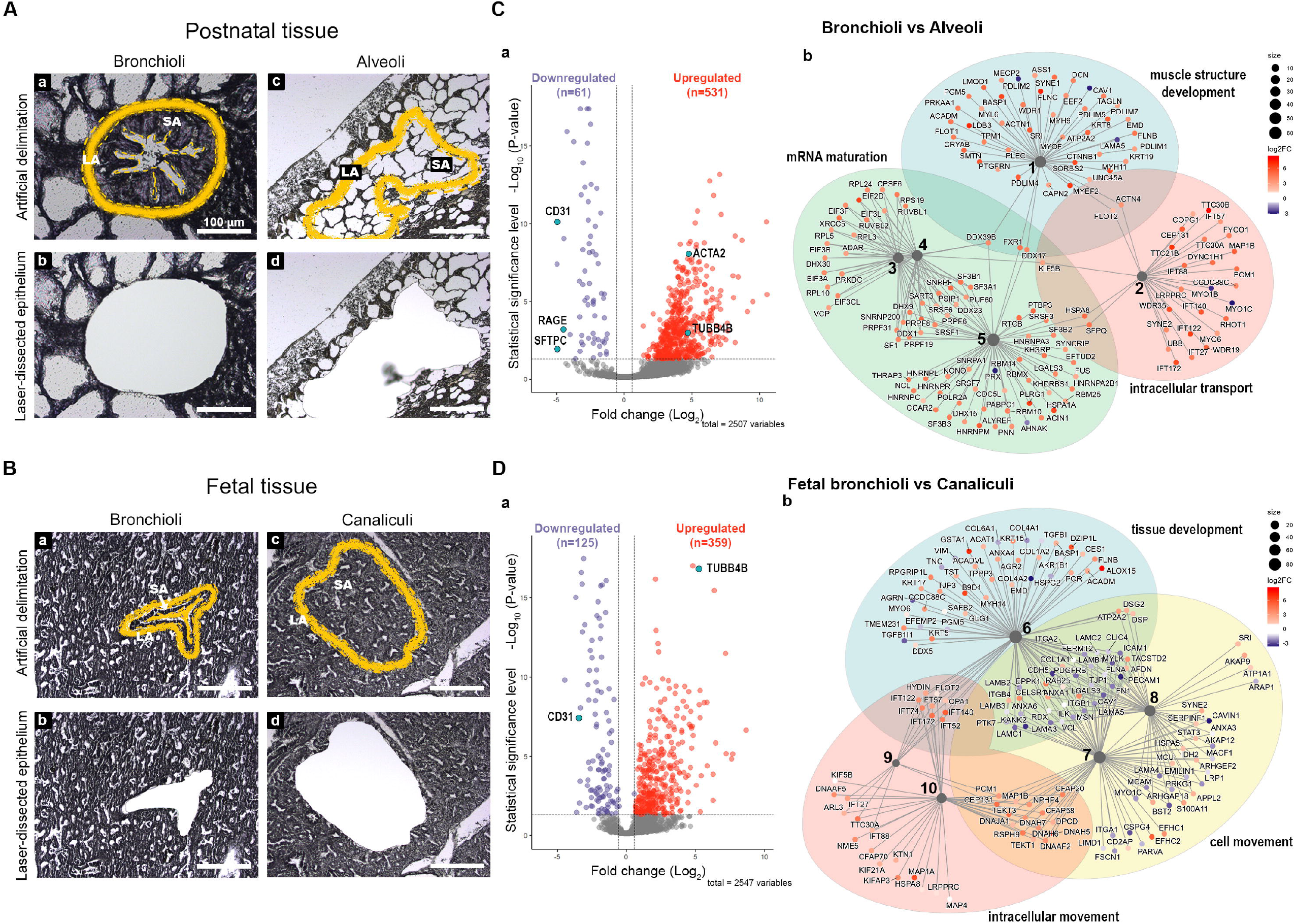
Differentially expressed proteins between postnatal bronchioli and alveoli, and fetal bronchioli and canaliculi. Scheme depicting the strategy for (A, a-d) postnatal and (B, a-d) fetal tissue LMD, depicting the selected area (SA) of tissue and the laser-destroyed area (LA). Volcano plots depicting the differential protein expression between (C, a) postnatal bronchioli and alveoli, and (D, a) fetal bronchioli and canaliculi, and their respective network plots (C, band D, b) of the enriched biological processes (BP). N=5 for each tissue structure. Proteins are identified by their gene name. BP legend: 1, muscle structure development; 2, cytoskeleton-dependent intracellular transport; 3, ribonucleoprotein complex assembly; 4, ribonucleoprotein complex subunit organization; 5, RNA splicing; 6, tissue development; 7, cell motility; 8, regulation of cellular component movement; 9, intraciliary transport; and 10, microtubule-based movement.

### 3.2 SIMPLE staining

To confirm the proteomic data, we performed SIMPLE staining, a technique developed by Glass and colleagues (7). Following the SIMPLE scheme depicted in **Figure 1**, fetal and postnatal lung samples (n=6 and n=9, respectively, with staining performed in triplicate for some of them) were stained for the different markers previously mentioned (**Figure 3**). For a proper validation of the technique, in addition to the tissue samples used for proteomic analysis, new samples were introduced (**Supplementary table 1**). The following order of primary antibodies was used: 1^st^ round staining CD31 and SFTPC, 2^nd^ round staining RAGE and ACTA2, and 3^rd^ round staining TUBB4B. For coverslip fixation we chose PBS since the use of Aquatex mounting solution was shown to affect, when using a primary antibody cocktail, the second antibody revelation. Aquatex was only used in the second revelation if a heat-induced antibody stripping was planned or if digital acquisition was not immediately possible. As expected and corroborating the proteomic data, RAGE, SFTPC and CD31 were detected in alveolar areas of postnatal tissue (**Figure 3A, panel a**), while they were virtually absent in fetal canaliculi **(Figure 3B, panel d**). Moreover, RAGE and SFTPC were restricted to postnatal alveoli and not present in other lung structures, while CD31 was evidently expressed by vascular endothelial cells of both postnatal (**Figure 3A, panel c)** and fetal tissues (**Figure 3B, panel f**). Regarding the bronchial regions, TUBB4B staining was detected in ciliary epithelial cells of both postnatal (**Figure 3A, panel b**) and fetal tissues (**Figure 3B, panel e**), with ACTA2 staining being specific for the airway smooth muscle layer surrounding the bronchioli. Alongside CD31, ACTA2 expression was expectedly present in vessels, staining vascular smooth muscle cells (**Figure 3A, panel c and 3B, panel f**). SOX-9, a well-known marker of chondrocyte differentiation, has been shown to play a central role in lung branching morphogenesis (16). Unexpectedly, SOX-9 was absent from the fetal data, which could be related either to a low abundance of this protein, illustrating a limitation of this proteomic pipeline, or to the age of the fetal samples. Indeed, these samples were of 19 and 21 gestational weeks (GW), and when performing IHC staining, SOX-9 was found at either low levels of expression or absent (**Supplementary figure 3**).To better characterize SOX-9 expression, we performed IHC staining, where SOX-9 was indeed detected at several early gestational timepoints (**Figure 4**), corroborating the results from others (16). In particular, we observed a strong expression of SOX-9 from our earliest time point, 12 gestational weeks (GW), until approximately 21 GW, where its expression decreased significantly (**Figure 4**). Noteworthy, at 14 and 16 GW, SOX-9 was found highly expressed (data not show). At later time points (26GW and postnatally) SOX-9 expression is, as expected, only detectable in cartilage tissue. To our surprise, in the course of our SIMPLE staining we observed that whenever SOX-9 was expressed, we were not able to detect RAGE expression. RAGE expression was only observed at later developmental stages, when SOX-9 expression was either low or absent (19-21 GW), gradually increasing until birth (26 GW-postnatal; **Figure 4**). To our knowledge this is the first description showing a timeline of RAGE protein expression in human fetal lung.

**Figure 3.**
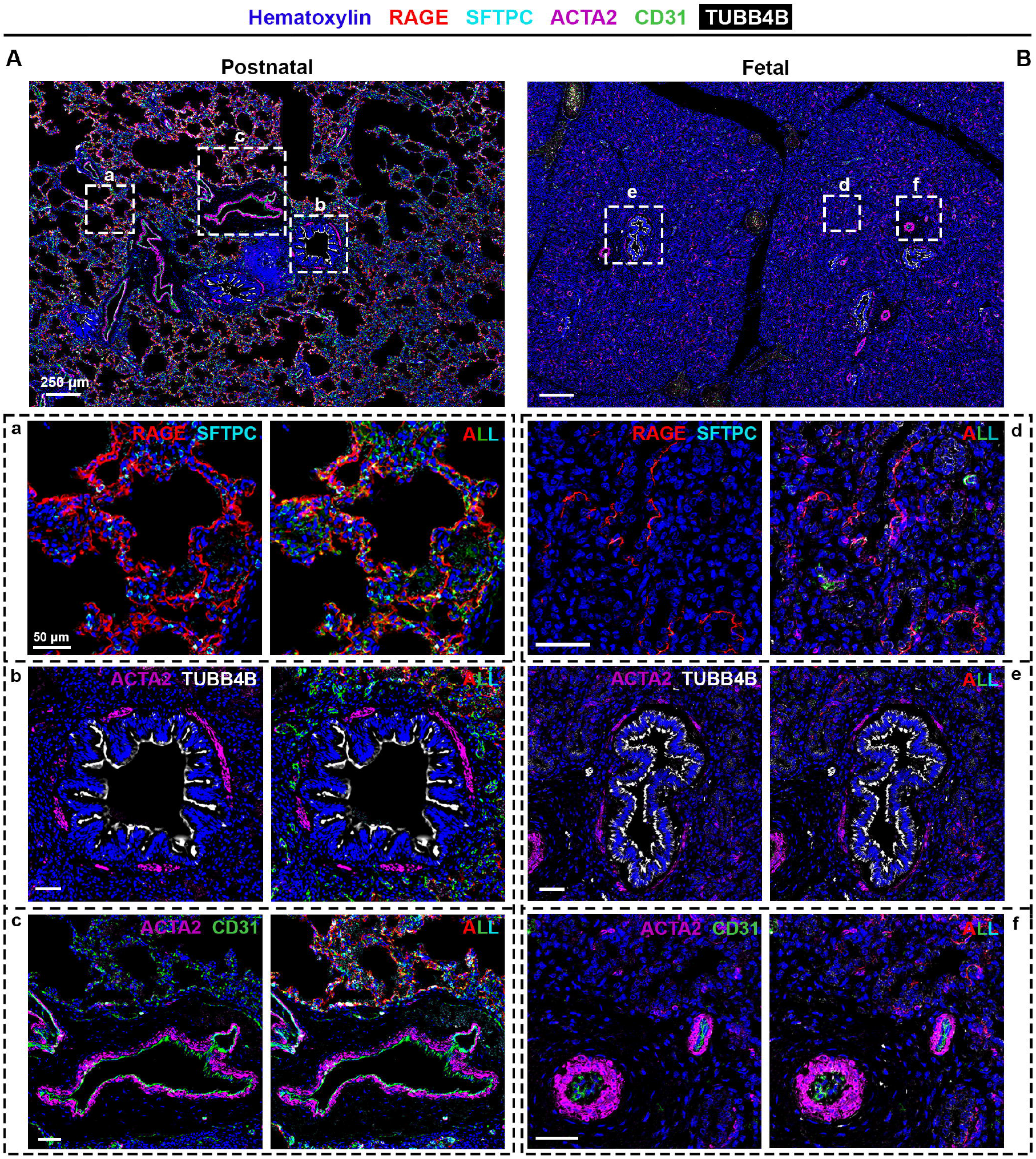
SIMPLE immunohistochemistry staining of **(A** and **a-c)** postnatal and **(B** and **d-f)** fetal human lung tissue. Postnatal and fetal lung tissues were stained against RAGE (red) and SFTPC (turquoise) to identify alveolar type I and type II pulmonary cells, typical of the **(a)** alveolar and **(d)** canalicular regions; TUBB4B (white) to identify ciliated cells, typical in **(b)** bronchioli and (**e)** fetal bronchioli; and ACTA2 (purple) and CD3 l (green), markers of smooth muscle cells and endothelial cells, respectively, present in (**c** and **f)** vessels. Hematoxylin is visualized in blue. N= 9 and n= 6 for postnatal and fetal tissue, respectively, with staining performed in triplicate for some of them.

**Figure 4.**
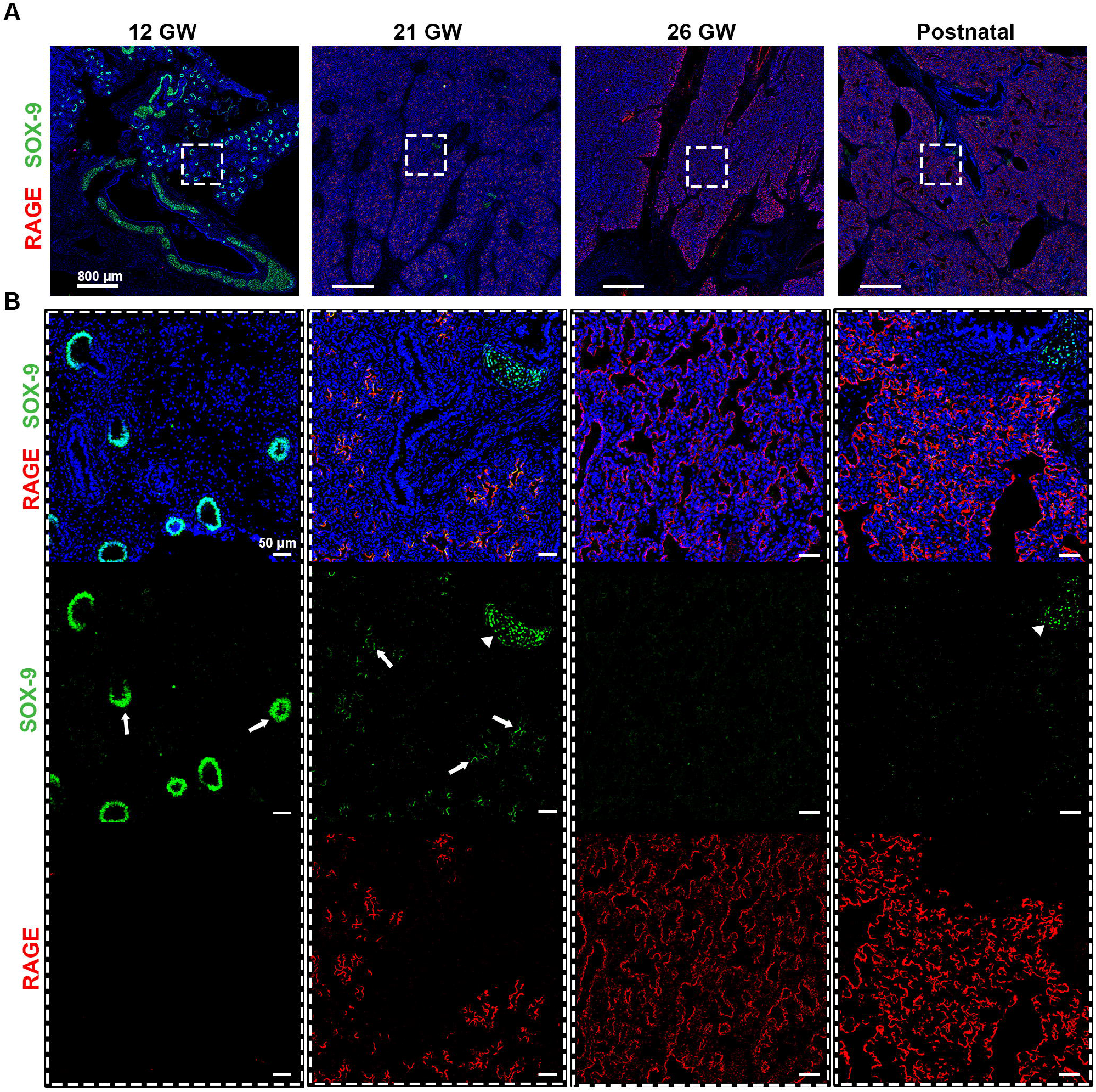
(A) Timeline for RAGE (red) and SOX-9 (green) expression in fetal tissues of 12, 21 and 26 gestational weeks (GW) and in postnatal tissue (10 days). (B) Magnification of the region of interest from the upper panel, where we can observe in separate, SOX-9 and RAGE. SOX-9 is expressed at earlier timepoints in canaliculi (arrows) and this expression is lost with the differentiation of airway cells. The loss of SOX9 is concumitant with the beginning of RAGE expression in early alveolar type I cells (in red). SOX-9 can also be found in the cartilage tissue (arrow heads). Hematoxylin is visualized in blue. N=3 for each timepoint of the fetal tissue. Staining was performed in replicate for some tissue.

## 4 Discussion

Human lung samples, especially to study rare developmental diseases, are difficult to obtain. In addition to the limited number of cases, collecting good quality retrospective samples of malformative or fetal lung cases is a real challenge. Fetal samples are also known to present preservation issues, due to the nature of the events that originate the collection of these tissues. In this report, we show that although the preservation process might not have been ideal, sample preparation and enrichment succeeded to create samples of satisfying quality and reproducibility, which allowed the subsequent comparative proteomic analysis. Nonetheless, we have decided to maintain samples of lower quality, such as sample E5 (**Supplementary figure 1**), to exemplify some of the caveats of this technique, which requires extensive handling in the preparation of the protein samples. In particular, sample E5 presents a high degree of missing values (represented in black, in Supplementary figure 1B) and this may have several origins, such as technical issues in LMD or sample preparation or tissue preservation. Interestingly, this sample shares the same source of tissue as sample G3 (alveoli), which also presents a high degree of missing values. Still on the alveolar samples (G1-G5), they presented the highest degree of variation intra-group, most probably due to the high number of missing values. In addition to the above-mentioned possible causes, differences in the honeycomb-like structure of alveoli might also affect the expected amount of obtained protein. Even though we took into consideration the suggested LMD areas, tissues may vary in structure, i.e. different proportions of cells to air space. Following the data quality validation, we corroborated our proteomic data by performing the SIMPLE technique, where we detected markers of bronchial and alveolar structures in healthy postnatal tissue and, when expected, in fetal tissue. This technique has been widely employed in both mouse and human tissues (7, 8), with most of these studies using tumor samples. In the work of Tsujikawa and colleagues, the authors performed an 11-round heat-mediated antibody stripping, which is mainly achievable due to the nature of the tissue. One of the main disadvantages of working with lung samples is its fragility, related to the honeycomb-like structure, in contrast to previous studies in the literature reporting results of the technique in more solid lung tumor samples. In this study we have empirically experienced that upon 5 rounds of HIER tissue degradation is quite extensive, which will certainly affect the specificity and reproducibility of the immunostaining. Other factors should be taken into considering for optimal immunostaining, such as: i) primary antibody specificity for species/tissues and cross-reaction of the secondary antibodies when using primary antibody cocktails; ii) antibody order, since not all epitopes present the same resistance to the chemical, mechanical and thermal manipulations during the staining process; and iii) balance between antibody stripping strength and tissue integrity, by testing the required conditions for a smooth, while proper, antibody removal.

We have also raised awareness for the limitations of a proteomic approach. As a proof of concept, we demonstrated that the transcription factor SOX-9, an important factor for cell differentiation, was not detected by proteomic, but was identified by IHC as being expressed early in development. The work of Danopoulos and colleagues corroborated our data, showing SOX-9 expression in fetal lung tissue during the pseudoglandular stage (4-16 weeks) until the canalicular stage (16-20 weeks), when SOX-9 expression starts decreasing (16). By performing IHC staining at different gestational timepoints, we observed that, in our samples, SOX-9 expression is lost between 19 and 21 GW. The variability might be related to the incertitude of the time of fecundation and its annotation, or simply due to the intrinsic biological variability. Therefore, the lack of SOX-9 signal in our proteomic data was most probably due to either a below the threshold level or absence of expression. In addition to a low basal level of protein expression that would not allow its proper detection, other possibilities could be related to the expected low amount of extracted protein from FFPE tissue or to the fact that nuclear proteins usually not as easily extracted/detected as cytoplasmic ones. Importantly, together with SOX-9 timeline, we have demonstrated, to the best of our knowledge for the first time, the timeline protein expression of RAGE. Its expression is inversely correlated with SOX-9 expression. Interestingly, our work complements the one of Clair and colleagues where they characterized RAGE expression, by LC-MS-based proteomic and Western blot, at 10 postnatal timepoints, from postnatal day 1 to 8 years (5). Finally, this work came as a technical proof of concept for LMD-based proteomic analysis completed by a SIMPLE multiplex staining, that will allow us to apply this approach in other contexts, such as in lung developmental diseases (e.g. congenital pulmonary airways malformations) where tissue accessibility and preservation are not optimal and alternatives not available.

Overall, and notwithstanding the intrinsic technical limitations of our pipeline, the results confirm the applicability and reproducibility of proteome-based analysis in FFPE pediatric lung tissues. We showed, by hand-picking specific markers, the differences between distinct lung structures (bronchioli, alveoli and vessels) and developmental state (fetal and postnatal tissue). Moreover, we have shown that proteomic data should be confirmed by other approaches, such as classical immunostaining, especially in the case of proteins of difficult extraction or low abundance.

## 5 Acknowledgements

This work was supported by the Ligue Pulmonaire Genevoise, the Fondation Privée de HUG and the Fondation Prim’Enfance. We thank the Proteomics and Bioimaging core facilities of the CMU, University of Geneva, for their equipment, advice and help. In particular, we thank Nicolas Liaudet from the Bioimaging core facility for his help in the development of the proper software for image processing, and Dre Labidi-Galy, from the department of medicine of the Hôpitaux Universitaires de Genève, for providing the necessary technical support related to the SIMPLE protocol.

