## Supplementary figures and images for "Laser microdissection, proteomics and multiplex immunohistochemistry: a bumpy ride into the study of paraffin-embedded fetal and pediatric lung tissues"

### Supplementary figure 1

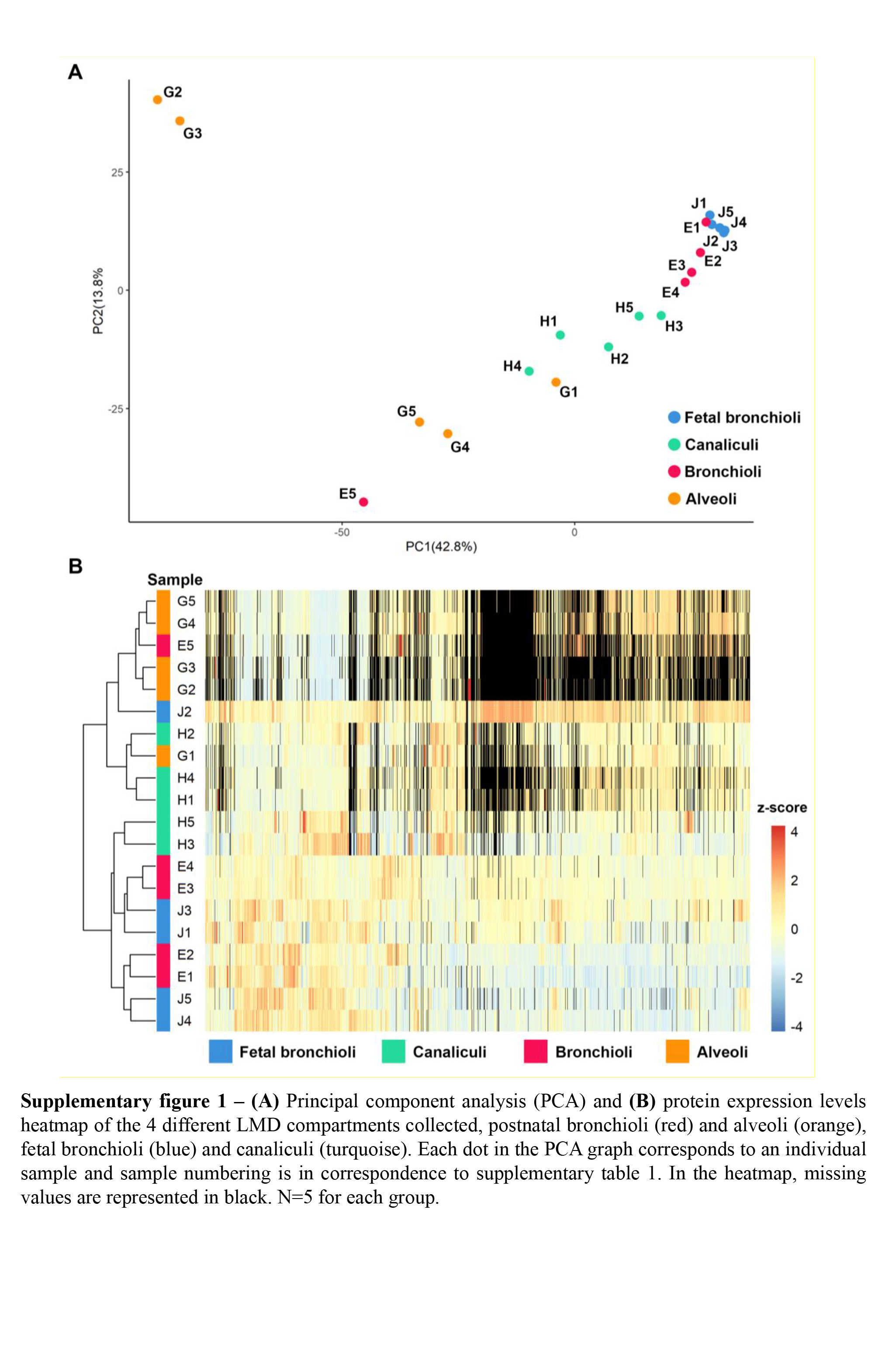

### Supplementary figure 2

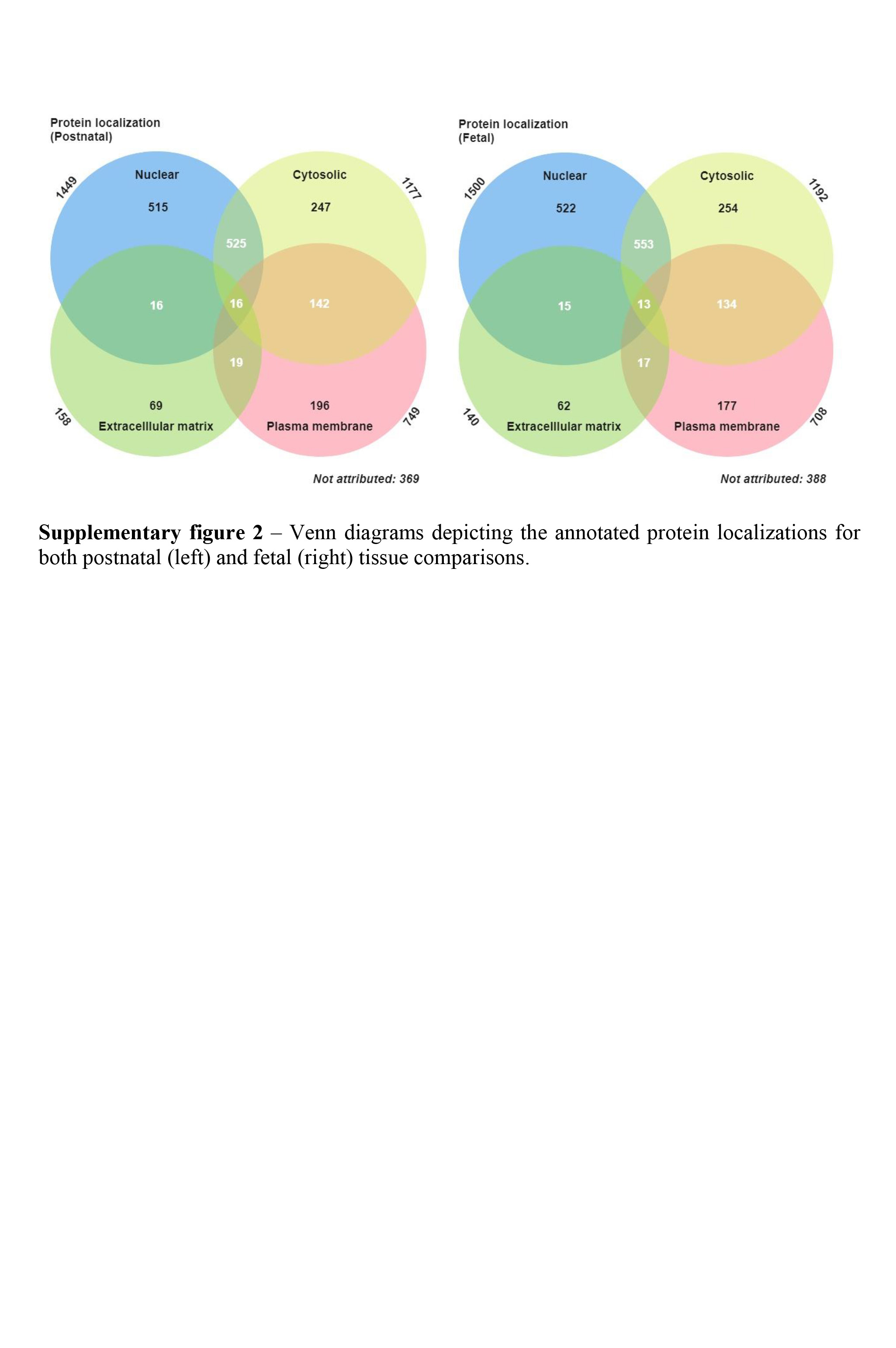

### Supplementary figure 3

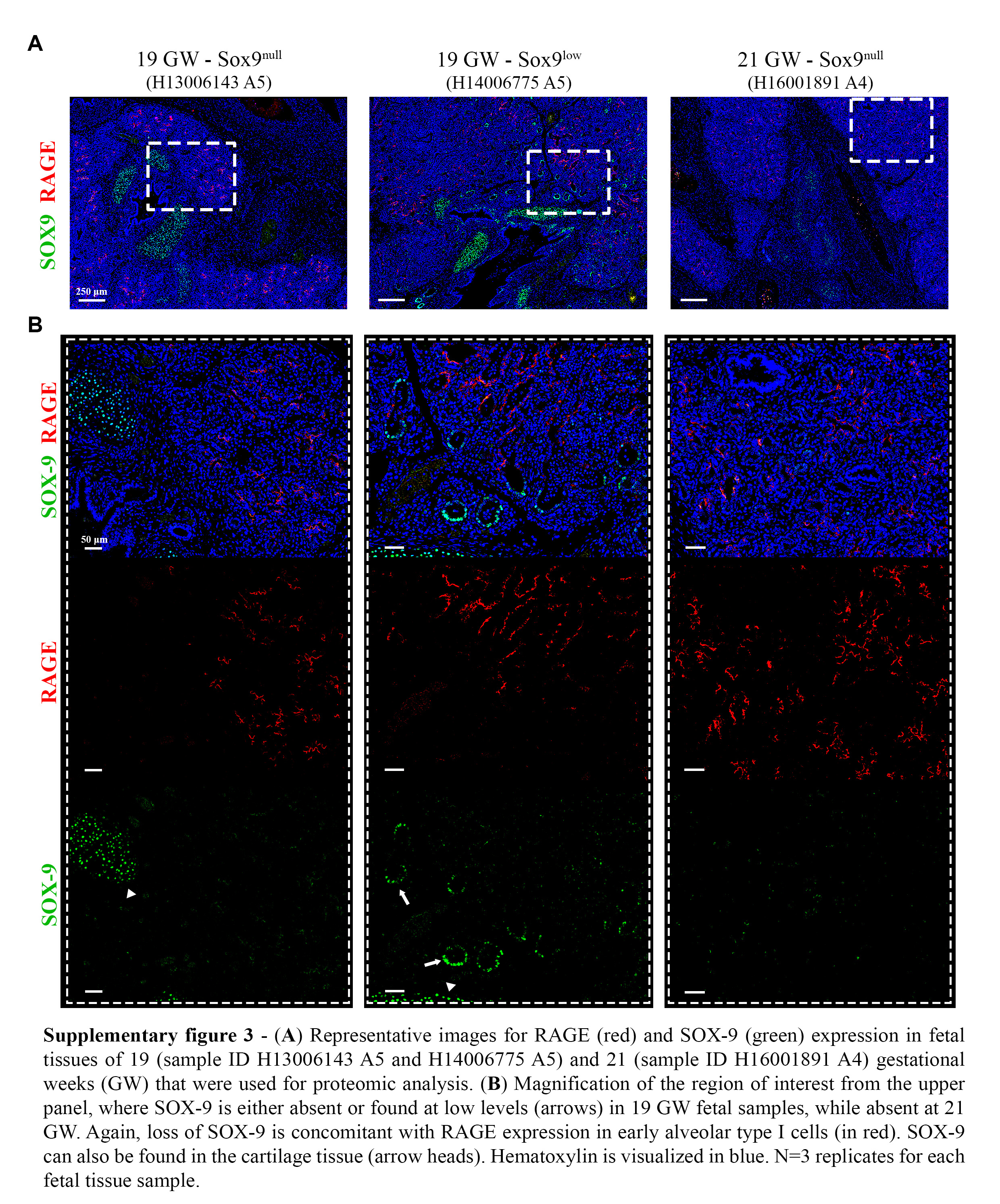
